# Derivation of endometrial gland organoids from term post-partum placenta

**DOI:** 10.1101/753780

**Authors:** Mirna Marinić, Vincent J. Lynch

## Abstract

A major limitation of recently developed methods for the generation of endometrial gland organoids is their reliance on decidua isolated from endometrial biopsies or elective termination of pregnancies. Here we report the successful establishment of endometrial gland organoids from decidua isolated from term placental membranes, as well as the identification of potential survival factors for the co-culture of gland organoids and endometrial stromal fibroblasts. These modifications facilitate the generation of patient-specific endometrial gland organoids with known pregnancy outcomes, such as term vs. preterm birth.

---

Determining the molecular mechanisms that underlie the establishment, maintenance, and cessation of pregnancy in humans is essential for understanding the basic biology of female reproduction as well as associated pathologies such as implantation failure, recurrent spontaneous abortion, and preterm birth. There are, however, significant technical and ethical limitations that prevent *in vivo* studies of human pregnancy. While animal models can overcome some of these limitations and be used to explore medically relevant questions, these systems do not recapitulate female reproductive traits that are unique to primates, apes, and humans such as extremely invasive hemochorial placentas, a derived (unknown) parturition signal and menstruation (Wildman *et al.*, 2006; Emera, Romero and Wagner, 2012; Wagner *et al.*, 2012). Unfortunately, the numerous *in vitro* models that have been developed since the 1980s to study pregnancy in humans are also limited because they rely on immortalized cells, primary cells with limited lifespans, single- or simple multi-cell culture that do not recapitulate the complex environment of the (pregnant) uterus, including interactions between maternal and fetal cell types (Lindenberg, Nielsen and Lenz, 1985; Weimar *et al.*, 2013 and references therein).

A promising solution to these problems is the development of organoid models of the maternal-fetal interface. Recently, Turco *et al.* (2017) and Boretto *et al.* (2017) demonstrated the derivation of endometrial gland organoids from human samples collected from endometrial biopsies and first trimester decidua. While the creation of methods to generate organoids from early pregnancy stages is important, biopsy and elective abortion samples can be difficult to obtain, are potentially fraught with ethical considerations, and have unknown pregnancy outcomes (e.g. term vs. preterm), thus cannot be used to generate patient-specific organoids with known pregnancy histories. To determine if endometrial gland organoids can be derived from post-partum tissue, we isolated decidua from term placental membranes and followed the protocol described in Burton *et al.* (2017) to derive gland organoids. Briefly, after digesting the decidua and releasing the gland elements from the surrounding stromal cells, glands were resuspended in Matrigel, plated in 48-well plates and overlaid with the Expansion Medium (ExM; **Supplementary Data**). Spheroids resembling previously described gland organoids (Turco *et al.*, 2017) began to appear 10-14 days after plating and could be passaged and recovered after freezing (Figure 1).

**Figure 1.**
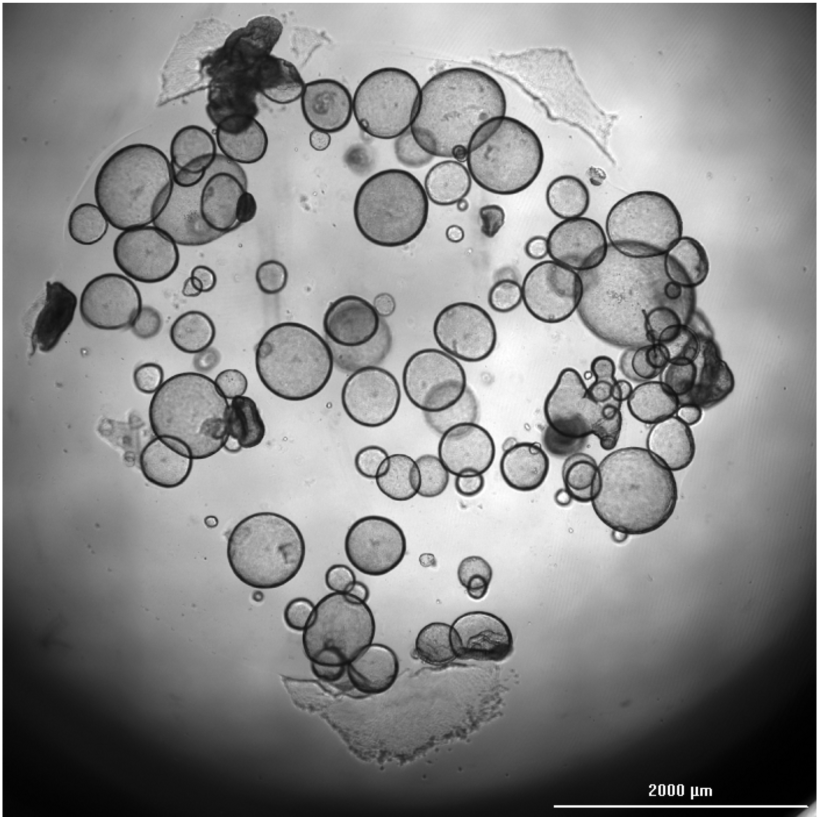
Endometrial gland organoids established from post-partum decidua sample. Scale bar, 2000μm. Image captured on Cytation-5 Cell Imaging Multi-Mode Reader (BioTek Instruments, Inc.).

We found that these organoids expressed classical markers of endometrial epithelial cells, including E-cadherin (E-CAD), Laminin (LAMA4), and Cytokeratin 7 (KRT7), consistent with an epithelial origin (Czernobilsky *et al.*, 1984; Aplin, Charlton and Ayad, 1988; Inoue *et al.*, 1992), as well as the progesterone receptor (PR), which is expressed by luminal and gland epithelia in the endometrium and decidua (Wang *et al.*, 1998) (Figure 2, Supplementary Figure 1). These organoids also express Vimentin (VIM; Figure 2), which is generally considered a marker of endometrial stromal fibroblasts (ESFs) (Can, Tekeliioǧlu and Baltaci, 1995), but is also present in endometrial glands *in vivo* (Dabbs, Geisinger and Norris, 1986; Matthews *et al.*, 1992). RNA-Seq of unprocessed glands, stromal cells, and organoids confirmed the expression of Vimentin in all three samples. Surprisingly, organoids clustered with endometrial stromal fibroblasts, rather than with isolated gland elements (Figure 3A). However, isolated glands likely contain multiple cell types giving them a unique gene expression profile. Indeed, numerous genes with expression enriched in the placenta are expressed in the gland samples, suggesting they contain placenta-derived cells (Table 1).

**Table 1.**
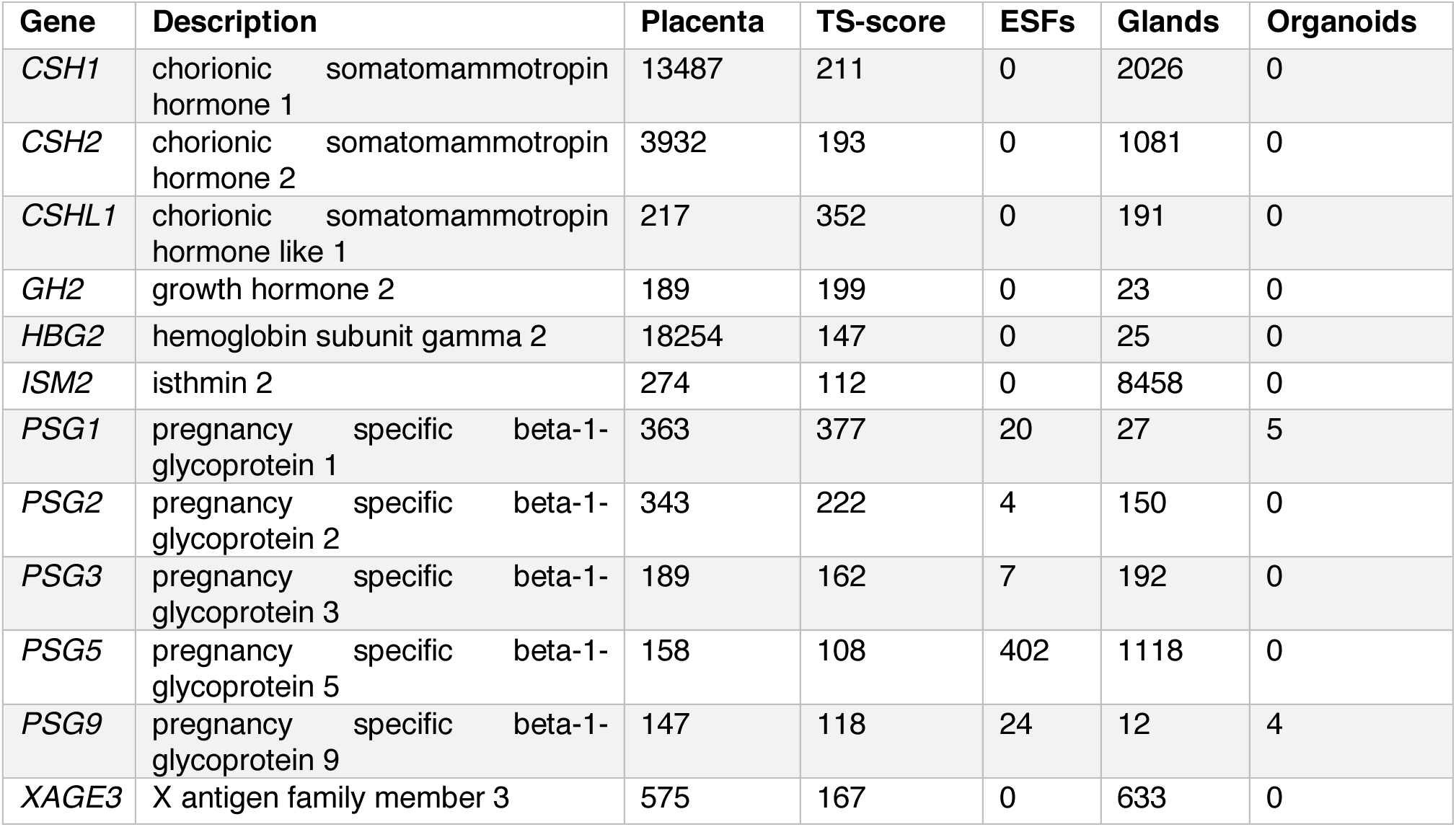
Expression of placental enriched genes in RNA-Seq data from placenta, endometrial stromal fibroblasts (ESFs), isolated glands before culture, and organoids after 4-8 passages. Expression levels are shown as transcripts per million (TPM) values, the Tissue Specificity score (TS-score) is calculated as the fold enrichment of each gene relative to the tissue with the second highest expression of that gene. TPM values are shown as averages of four replicates for ESFs, and three replicates for glands and organoids. Placental data is from: https://www.proteinatlas.org/humanproteome/tissue/placenta.

**Figure 2.**
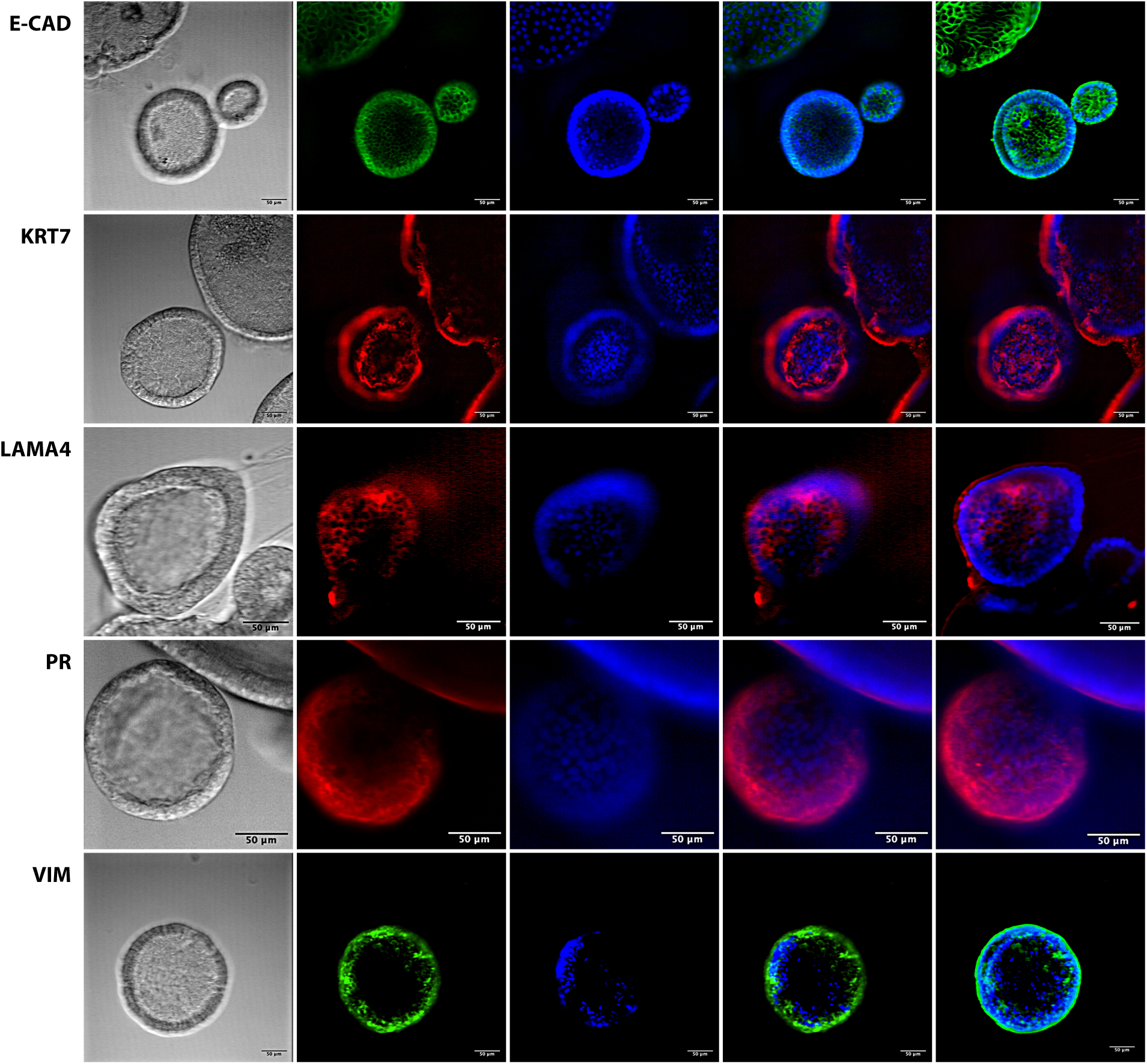
Immunofluorescent staining of the endometrial gland organoids. Proteins detected: E-cadherin (E-CAD); Cytokeratin 7 (KRT7); Laminin (LAMA4); Progesterone receptor (PR); Vimentin (VIM). Panels show (left to right): bright field; detected protein; DAPI-stained nuclei; merge at the level of 1 selected slice; flattened Z-stack merge of all slices. Scale bars, 50μm.

**Figure 3.**
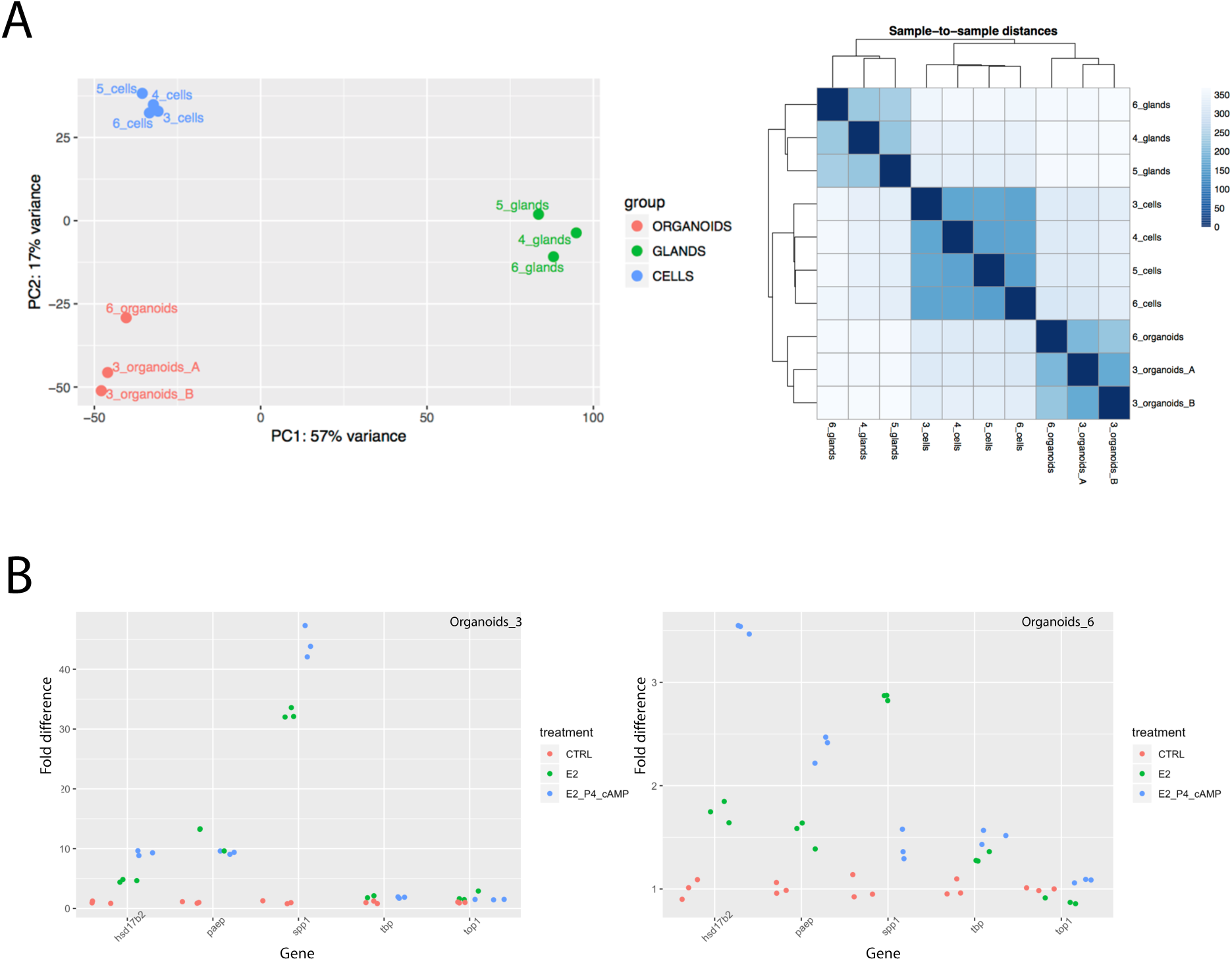
RNA-Seq and qRT-PCR results. A, PCA plot and Sample-to-Sample Distance graph of RNA-Seq data from ESFs, unprocessed glands, and organoids (donors 3, 4, 5 and 6). B, qRT-PCR data for hormonal treatment of organoids. Treatment was performed on samples derived from 2 different donors (3 and 6). Tested genes: *PAEP*, *SPP1* and *HSD17B2*. Control housekeeping genes: *TBP* and *TOP1*.

We next sought to determine if these gland organoids were hormone responsive by treating them with β-estradiol (E2), progesterone (P4) and adenosine 3’,5’-cyclic monophosphate (cAMP), and assaying gene expression by qRT-PCR. We verified that *PAEP*, *SPP1* and *HSD17B2*, previously characterized markers of hormone responsiveness in glands and gland organoids (Borthwick *et al.*, 2003; Kamarainen *et al.*, 1993; Briese *et al.*, 2006; Satyaswaroop, Wartell and Mortel, 1982; Turco *et al.*, 2017), were differentially expressed after hormone treatment (Figure 3B). These data indicate that gland organoids derived from term placentas maintain appropriate hormone responsiveness after isolation and culture. Collectively, our results demonstrate that endometrial gland organoids can be established from decidua isolated from term placentas, allowing for the generation of patient-specific organoids with known pregnancy outcomes.

To establish an organoid model that better recapitulates the cellular composition of the endometrium, we also isolated endometrial stromal fibroblasts (ESFs) from placentas for co-culture with organoids. We found that ESFs cultured in ESF media grew well, whereas they died within a few days when cultured in ExM. Surprisingly, however, ESFs survived in ExM initially conditioned by organoids for three days, suggesting that organoids express an ESF survival factor(s). This prompted us to analyze organoid-conditioned ExM by liquid chromatography coupled with tandem mass spectrometry (LC-MS/MS) to identify secreted proteins that may function in promoting ESF survival in ExM. We found 11 proteins that were significantly enriched in organoid-conditioned ExM compared to control ExM, including 5 that are not expressed by ESFs (**Supplementary Table 1**). While further investigation of these findings is needed, our results suggest these secreted proteins act as possible survival factors.

Here we demonstrate that endometrial gland organoids can be derived from decidua isolated from post-partum placentas, facilitating the establishment of patient-derived organoids, while providing greater simplicity and minimizing ethical objections associated with tissues collected from biopsies or elective pregnancy terminations. Our observation that ESFs can survive in ExM also opens the possibility of building a more realistic 3-dimensional representation of maternal-fetal interface *in vitro*, incorporating multiple biologically relevant cell types.

## Supporting information

Supplementary Dats

## Author contributions

M.M. performed all the experiments, analyzed the data and wrote the manuscript.

V.J.L. supervised the project, analyzed the data and wrote the manuscript.

## Acknowledgements

We are grateful to all the women who donated placental tissue samples, without which this research would not have been possible. We would also like to thank Dr. Sarosh Rana, MD, and the University of Chicago Medicine hospital team who collected the placental samples; Cristina Paz and the March Of Dimes Membrane Processing Core in the Human Genetics Department of the University of Chicago for the initial processing of the samples and providing the isolated decidua; Dr. Margherita Y. Turco for kind input on the details of the original protocol.

## SUPPLEMENTARY DATA

### Establishment of Endometrial Gland Organoid Culture

Placental samples were collected from women delivering at term at the University of Chicago Medicine hospital, following written consent. All post-partum placental tissue samples measuring ~5×5cm were collected in RPMI 1640 [Thermo Fisher Scientific, 11875093] with 1x Penicillin-Streptomycin [Thermo Fisher Scientific, 15140122]. Initial processing (separating amnion, trophoblast and decidua) was done within 24 hours upon receiving at the March of Dimes Membrane Processing Core in the Human Genetics Department of the University of Chicago.

Isolated decidua were treated as described in Burton *et al.* (2017). In short, decidua were minced in a dish, followed by digestion in collagenase V (0.4mg/ml)/dispase II (1.25U/ml) solution [Sigma, C9263 and D4693] in RPMI-1640 with 10% Fetal Bovine Serum [FBS; Thermo Fisher Scientific, 26140-079] for at least 20 minutes at 37°C with shaking. The digestion length is critical, in order for the stromal cells to be separated from gland elements, but without over-digesting the glands. Presence of significant mucous and greater thickness of the tissue samples usually requires longer digestion time. The digestion was stopped by dilution with RPMI-1640 and the digest was filtered through several 100μm sieves. After rinsing the sieves with RPMI-1640, gland elements were collected by inverting the sieves over the Petri dish and washing them off. They were pooled into low binding tubes [Eppendorf, 022431081] and pelleted by centrifugation at 600g for 6 minutes, re-suspended in Advanced DMEM/F12 [Thermo Fisher Scientific, 12634010] and pelleted again. After removing the supernatant, gland elements were placed on ice and mixed with 20x volume of Matrigel [Corning, 356231]. 20-25μl droplets were plated per well of a 48-well plate [Thomas Scientific, 1156F01]. They were put in the humidified cell culture incubator at 37°C and 5% CO_2_ to solidify for 15 minutes, overlaid with 250μl of Expansion Medium (ExM, see below) and returned to the incubator. ExM was exchanged every 3 days. First spherical structures start appearing about 10-14 days post-plating. After subsequent passaging, it takes them about 2-3 days to appear.

Expansion Medium (ExM) composition, as in Burton *et al.* (2017):

Advanced DMEM/F12 [Thermo Fisher Scientific, 12634010]

L-glutamine, 2mM [Thermo Fisher Scientific, 25030-081]

B27 minus vitamin A supplement, 1x final [Thermo Fisher Scientific, 12587010] N2 supplement, 1x final [Thermo Fisher Scientific, 17502048]

Nicotinamide, 10nM [Sigma, N0636]

N-acetyl-L-cysteine, 1.25mM [Sigma, A9165]

Primocin 100μg/ml [Invivogen, ant-pm-1]

ALK-4, −5, −7 inhibitor, 500nM [StemRD, A8-002]

Recombinant human R-spondin-1, 500ng/ml [Peprotech, 120-38]

Recombinant human Noggin, 100ng/ml [Peprotech, 120-10C]

Recombinant human FGF-10, 100ng/ml [Peprotech, 100-26]

Recombinant human EGF, 50ng/ml [Peprotech, AF-100-15]

Recombinant human HGF, 50ng/ml [Peprotech, 100-39]

Y-27632, 10μM [Stemcell, 72302], added to ExM only for the first 3 days after thawing and re-plating the organoids

### Passaging and Freezing

Passaging and freezing were done as described in Burton *et al.* (2017). Passaging was performed as needed, when organoids were abundant and still spherical. They can grow large, but then they start collapsing onto themselves.

Pelleting was performed by centrifugation at 600g for 6 minutes and, unless otherwise noted, in 1ml of Advanced DMEM/F12.

#### Passaging

Matrigel droplets containing organoids were scraped off the plate using low binding pipette tips, transferred together with ExM to the low binding tubes (3-4 wells per tube) and pelleted. After removing the supernatant and adding 250μl of Advanced DMEM/F12, organoids were broken by pipetting up-and-down for 300 times. Broken organoids were pelleted again and pipetted up-and-down for additional 80 times. After final centrifugation and removing the supernatant, pellets were cooled on ice for 1-2 minutes, resuspended in Matrigel and 20-25μl droplets were plated as described above. For breaking the Matrigel droplets, we used Picus electronic pipette 50-1000ul [Sartorius, 735081]. First 300x pipetting step was performed at setting 9 (highest), second 80x pipetting step at setting 7.

#### Freezing

Before freezing, ExM was exchanged for 250μl Cell Recovery Solution [Corning, 354253] and the organoids were incubated for 1 hour on ice. Well content was collected in low binding tubes and pelleted. After removing the supernatant, organoids were washed with 1ml of Advanced DMEM/F12 and pelleted again. Finally, the pellet was resuspended in 500μl Recovery Cell Culture Freezing Medium [Thermo Fisher Scientific, 12648010], transferred to cryovials in a freezing container and left at −80°C overnight. Long-term storage should be in the vaporous phase of liquid N_2_.

### Establishment of Primary Endometrial Stromal Fibroblasts Culture

After stopping the decidua digest and filtering through 100μm sieve, 6ml aliquot from the flow-through was directly pipetted into T25 cell culture flask and placed in the humidified cell culture incubator at 37°C and 5% CO_2_. The next day, the remaining floating blood cells were washed away with DPBS [Thermo Fisher Scientific, 14190250] and the ESF medium was added (see below). Medium was exchanged every 3 days and the expansion of few initially attached endometrial stromal fibroblasts (ESFs) was observed. It takes about 2 weeks for the cells to become confluent. From then on, ESFs can be expanded by passaging every 2-3 days.

To passage ESFs, cells are rinsed with DPBS and de-attached from the dish by incubation in 0.05% Trypsin-EDTA, no phenol red [Thermo Fisher Scientific, 15400-054] for 3-5 minutes in the humidified incubator at 37°C and 5% CO_2_. Trypsinisation is stopped with the ESF medium and cells are pelleted by centrifugation at 200g for 5 minutes. Pellet is resuspended in the appropriate volume of medium and cells were plated to new cell culture flasks. For the long-term storage, cells can be frozen and kept in the vaporous phase of liquid N_2_. The freezing medium is ESF medium supplemented with 5% dimethyl sulfoxide [DMSO; Thermo Fisher Scientific, BP231].

ESF medium:

Phenol Red-free DMEM [Thermo Fisher Scientific, 31053-028]

10% FBS [Thermo Fisher Scientific, 26140-079],

1% L-glutamine [Thermo Fisher Scientific, 25030-081]

1% sodium pyruvate [Thermo Fisher Scientific, 11360070]

1x insulin-transferrin-selenium [ITS; Thermo Fisher Scientific, 41400045]

### Immunofluorescence and Confocal Microscopy

When organoids were abundant, they were fixed directly in 48-well plate as follows: ExM was removed and exchanged with 250μl cold 4% paraformaldehyde (PFA) in DPBS. Organoids were incubated for 20 minutes at room temperature and washed 3×10 minutes in 500μl DPBS prior to staining.

#### Staining steps

If not otherwise noted, all the steps were performed at room temperature and all the washes were done 3×10 minutes in 500μl DPBS. For detecting intra-cellular progesterone receptor (PR), fixed organoids were permeabilized for 10 minutes in 250μl 0.25% Triton X-100 in DPBS and washed. Then 500μl of blocking reagent (DPBST [DPBS + 0.1% Tween] + 1% Bovine Serum Albumin [BSA; ChemCruz, sc-2323] + 22.52mg/ml glycine [Sigma, 8.16013]) was added for 1 hour, followed by washing. Incubation with primary antibodies (diluted in 250μl DPBST+1% BSA) was done over night at 4°C. The next day organoids were washed and incubated with secondary antibodies (diluted in 250μl DPBST+1% BSA) for 1 hour. After washing, organoids were counterstained for 1 minute with 0.5-1μg/μl of DAPI [Thermo Fisher Scientific, 62247] in DPBS shielded from the light, washed one final time 3×5 minutes and were kept at 4°C in the dark until imaging.

| ANTIBODY | PROTEINTECH CAT. # | DILUTION |
| --- | --- | --- |
| **Primary** |  |  |
| E-cadherin<br>Mouse Monoclonal | 60335-1-Ig | 1:200 |
| Cytokeratin 7-specific<br>Rabbit Polyclonal | 17513-1-AP | 1:50 |
| LAMA4 (Isoform 3)<br>Rabbit Polyclonal | 10465-1-AP | 1:100 |
| Progesterone Receptor<br>Rabbit Polyclonal | 25871-1-AP | 1:20 |
| Vimentin<br>Mouse Monoclonal | 60330-1-Ig | 1:200<br>1:2000* |
| **Secondary** |  |  |
| Goat anti-mouse IgG<br>(H+L)<br>FITC conjugate | SA00003-1 | 1:200 |
| Goat anti-rabbit IgG (H+L)<br>R-PE conjugate | SA00008-2 | 1:75 |

\*Both dilutions gave matching results.

Images were captured on an Olympus IX81 microscope with a DSU spinning disk confocal unit [Olympus Corporation of the Americas, Center Valley, PA] using a Hamamatsu model C9100 EM-CCD camera [Hamamatsu Photonics, Skokie, IL] run by SlideBook v6.0 software [Intelligent Imaging Innovations, Denver, CO] at the Integrated Microscopy Core Facility at the University of Chicago. Image processing was done in ImageJ version 2.0.0-rc-69/1.52 and Adobe Photoshop CS4 version 34.0.0 [Adobe Systems Inc.].

### RNA Isolation and cDNA Synthesis

Organoids were incubated in 250μl Cell Recovery Solution for 1 hour on ice. Well content was collected in low binding tubes and pelleted by centrifugation at 600g for 6 minutes. After removing the supernatant, organoids were washed with 1ml of Advanced DMEM/F12 and pelleted again. RNA was isolated from the pellet using RNeasy Plus Mini kit [QIAGEN, 74134] following manufacturer’s protocol.

cDNA was synthesized from 200ng input RNA, using the Maxima H Minus First Strand cDNA Synthesis Kit [Thermo Fisher Scientific, K1652] with oligo(dT)_18_ primers and incubating samples as follows: 10 minutes at 23°C, 15 minutes at 55°C, 5 minutes at 80°C.

### Hormonal Treatment and qRT-PCR

Organoids from 2 different donors were passaged and grown for 2-4 days, treated with 10nM β-estradiol [E2, Sigma E4389] in ExM for 48 hours and then treated with 10nM β-estradiol, 1μM progesterone [P4, Sigma P7556] and 1μM 8-bromoadenosine 3′,5′-cyclic monophosphate [cAMP; Sigma B5386]. After an additional 48 hours, RNA was isolated as described above and relative expression levels of *PAEP* (Hs01046125_m1), *SPP1* (Hs00959010_m1) and *HSD17B2* (Hs00157993_m1) were measured in triplicates by qRT-PCR. Control housekeeping genes were *TBP* (Hs00427620_m1) and *TOP1* (Hs00243257_m1). We used TaqMan Fast Universal PCR Master Mix 2x [Thermo Fisher Scientific, 4352042] and followed manufacturer’s protocol. DeltaCt values were calculated using Microsoft Excel and fold differences were plotted as a scatter plot in RStudio version 1.2.1335 [RStudio, Inc.].

qRT-PCR program:

1. 95°C, 30s
2. 95°C, 5s
3. 60°C, 30s + plate read
4. GO TO 2 x 50
5. END

### RNA-Seq and Data Analysis

RNA was isolated from: unprocessed gland elements (small aliquot collected from the sieve after filtering the initial digest of decidua), growing organoids and endometrial stromal fibroblasts isolated from the same samples and established as primary cell cultures. After DNase treatment with TURBO DNA-free Kit [Thermo Fisher Scientific, AM1907], RNA quality and quantity were assessed on 2100 Bioanalyzer [Agilent Technologies, Inc.]. RNA-Seq libraries were prepared using TruSeq Stranded Total RNA Library Prep Kit with Ribo-Zero Human [RS-122-2201, Illumina Inc.] and following manufacturer’s protocol. Library quality and quantity were checked on 2100 Bioanalyzer and the pool of libraries was sequenced on Illumina HiSEQ4000 (single-end 50bp) using manufacturer’s reagents and protocols. DNase treatment, quality control, Ribo-Zero library preparation and Illumina sequencing were performed at the Genomics Facility at The University of Chicago.

All sequencing data were uploaded and analyzed on the Galaxy platform (public server https://usegalaxy.org, Version 19.05), converted using FASTQ Groomer and aligned using HISAT2. We assembled transcripts using StringTie, with reference GTF file and “Reference Transcripts Only” option, followed by DESeq2 to calculate differential gene expression. The reference GTF file for guided assembly was done by StringTie merge of separately generated String Tie files (this time without “Reference Transcripts Only” option) and ensGene table from “Ensembl Genes” track from UCSC Genome Browser (GRC37/hg19 assembly).

### Mass Spectrometry Analysis of Conditioned ExM

Fresh ExM was added to the growing organoids and also to the empty Matrigel droplets. After 3 days in the culture, organoid-conditioned and Matrigel-conditioned media were collected, as well as the aliquot from unconditioned ExM. 1x of Halt^TM^ Protease Inhibitor Single-Use Cocktail 100x [Thermo Fisher Scientific, 78430] was added to all 3 samples and the samples were snap-frozen in liquid N_2_.

Sample preparation, LC-MS/MS and data analysis were performed at the Mass Spectrometry Core of The University of Illinois at Chicago as described below.

#### Sample preparation

100μl of each sample was taken and processed following standard S-Trap protocol [Protifi, S-Trap™ micro kit, K02-micro-10]. Samples underwent tryptic digestion and desalting, the residue was reconstituted into 50μl of 5% ACN in water with 0.1% FA, then further diluted 10-fold for LC-MS/MS analysis.

#### LC-MS/MS method

Samples were analyzed by nanoLC-HRMS system [Thermo Orbitrap Velos Pro]. For each sample 1μl was injected, peptides were eluted on a reverse phase C18 column via 1 hr gradient LC program. Mobile phase A: water with 0.1% FA; mobile phase B: ACN with 0.1% FA. Data were acquired by a DDA method (data dependent acquisition: 1 full scan plus 12 MS2 scan for the most intense 12 ions). Raw data were submitted to Mascot and searched against UniProt human proteome database. Search results were validated by Scaffold [version 4.9.0; Proteome Software Inc.]. Search parameters can be found in the Scaffold result file in the section of “Publish”. For the original result file, see **Supplementary Data File 1**.

## WEB RESOURCES

### UCSC Genome Browser

https://genome.ucsc.edu/

### ENSEMBL Genome Browser and BioMart tool

http://useast.ensembl.org/index.html

http://useast.ensembl.org/biomart/martview/

### The Human Protein Atlas

https://www.proteinatlas.org/

### Galaxy version: 19.05 (Afgan *et al.*, 2018)

https://usegalaxy.org

### Galaxy tools

**FASTQ Groomer**, converts between various FASTQ quality formats, Galaxy Version 1.1.1 (Blankenberg *et al.*, 2010)

**HISAT2**, a fast and sensitive alignment program, Galaxy Version 2.1.0+galaxy4 (Kim, Langmead and Salzberg, 2015)

**StringTie** transcript assembly and quantification and **StringTie merge** transcripts, Galaxy Version 1.3.4 (Pertea *et al.*, 2015)

**DESeq2**, determines differentially expressed features from count tables, Galaxy Version 2.11.40.2 (Love, Huber and Anders, 2014)

---

**Supplementary Figure 1.**
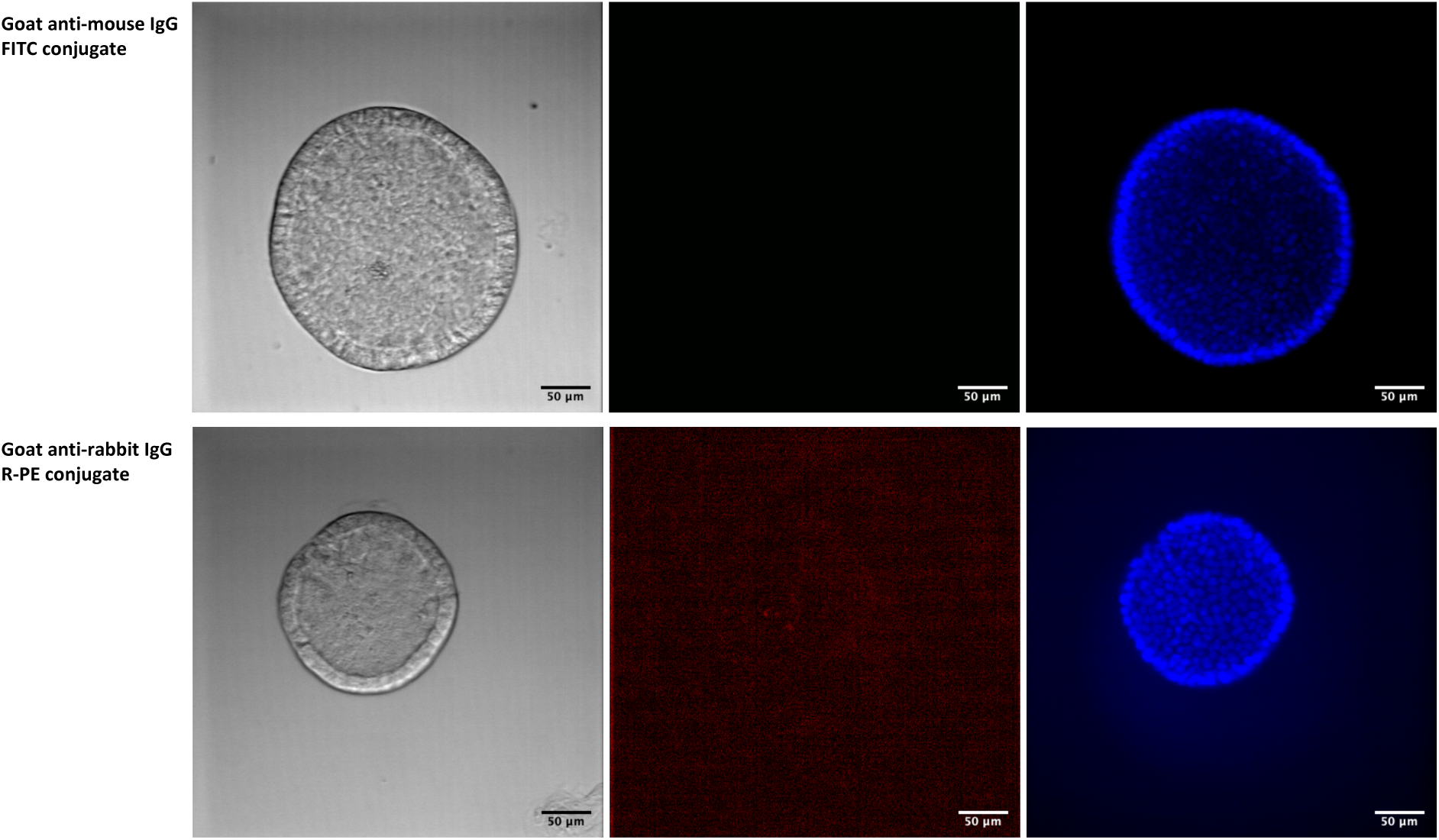
Control immunofluorescent staining of the endometrial gland organoids. Organoids were incubated only with secondary antibodies to test for the background staining. Antibodies used: goat anti-mouse IgG (H+L) FITC conjugate (green) and goat anti-rabbit IgG (H+L) R-PE conjugate (red). Scale bar: 50μm.

**Supplementary Table 1.** Mass spectrometry results of conditioned ExM. Orange highlighted rows show proteins uniquely detected in organoid-conditioned ExM. Green highlighted proteins are found to be expressed in endometrial and cervical glands, but not in the stromal cells (source: Human Protein Atlas). FDR=1% for protein and peptide threshold.

**Supplementary Data File 1.** Original mass spec data.

